# M2 Cortex-Dorsolateral striatum stimulation reverses motor symptoms and synaptic deficits in Huntington’s Disease

**DOI:** 10.1101/2020.04.08.032359

**Authors:** Sara Fernández-García, Sara Conde-Berriozabal, Esther García-García, Clara Gort-Paniello, David Bernal-Casas, Gerardo García-Díaz Barriga, Javier López-Gil, Emma Muñoz-Moreno, Guadalupe Sòria, Leticia Campa, Francesc Artigas, Manuel J Rodriguez, Jordi Alberch, Mercè Masana

## Abstract

Huntington’s disease (HD) is a neurological disorder characterized by motor disturbances. HD pathology is most prominent in the striatum, the central hub of basal ganglia. The cortex is the main striatal afference and progressive cortico-striatal disconnection characterizes HD. We mapped cortico-striatal dysfunction in HD mice to ultimately modulate the activity of selected cortico-striatal circuits to ameliorate motor symptoms and recover synaptic plasticity. Multimodal MRI *in vivo* suggested prominent functional network deficits in fronto-striatal compared to motor-striatal pathways, which were accompanied by reduced glutamate levels in the striatum of HD mice. Moreover, optogenetically-stimulated glutamate release from fronto-striatal terminals was reduced in HD mice and electrophysiological responses in striatal neurons were blunted. Remarkably, repeated M2 Cortex-dorsolateral striatum optogenetic stimulation normalized motor behavior in HD mice and evoked a sustained increase of synaptic plasticity. Overall, these results reveal that the selective stimulation of fronto-striatal pathways can become an effective therapeutic strategy in HD.

## Introduction

Huntington’s disease (HD) is an inherited neurodegenerative disorder with symptomatic manifestations including involuntary movements such as chorea, dystonia, poor motor coordination, psychiatric and cognitive symptoms. HD is caused by a CAG repeat expansion in the huntingtin gene (Htt) that translates into a polyglutamine tract in the huntingtin protein. Mutant Htt (mHtt) is expressed throughout the brain and disrupts a wide range of molecular pathways and signaling cascades, yet HD pathology is most prominent in the basal ganglia circuitry.

The main hub of the basal ganglia is the striatum, which controls movements and behavior through a myriad of inputs and outputs (Calabresi et al., 2014; Freeze et al., 2013; Rothwell et al., 2015; Rueda-Orozco and Robbe, 2015; Tecuapetla et al., 2016). The striatum receives glutamatergic inputs from all neocortical areas and routes the information through the basal ganglia (Bolam et al., 2000). Information flows from the cortex through the basal ganglia and goes back to the cortex via the thalamus through two main pathways (direct and indirect), which orchestrate the proper execution of movement. Additionally, the specific origin of the cortical inputs to the striatum provides a substrate for information segregation in these circuits (Hintiryan et al., 2016; Wall et al., 2013).

In HD, motor symptoms emerge from dysregulated information flow through the basal ganglia circuits. In particular, disturbances of the cortico-striatal communication seem to have a leading role in HD network dysfunction, with alterations appearing in prodromal HD (Burgold et al., 2019; Dumas et al., 2013; Unschuld et al., 2012). Specifically, the caudate nucleus and the premotor cortex appear to be predominantly affected (Unschuld et al., 2012). In HD mouse models, several evidences point to a progressive disconnection between cortex and striatum (Cepeda et al., 2007; Veldman and Yang, 2018). Cortico-striatal dysfunction in HD is strongly supported by the impaired cortico-striatal-dependent motor functions (Hong et al., 2012; Puigdellívol et al., 2015), altered paired-pulse facilitation at cortico-striatal synapses (Milnerwood and Raymond, 2007), reduction of striatal excitatory postsynaptic currents with loss of cortico-striatal synapses (Cepeda et al., 2003; Deng et al., 2013); and altered glutamate release in striatum of transgenic R6/1 mice (NicNiocaill et al., 2001). Moreover, striatal pathology is reduced in mouse lines by restricting mHtt expression to striatum, as opposed to striatum and cortex (Gray et al., 2008; Gu et al., 2005; Wang et al., 2014). Thus, a progressive disconnection of cortico-striatal pathways alters the information processing in basal ganglia circuitry, leading to motor disturbances. However, we do not know yet which specific cortico-striatal pathways are affected.

In this study, we aimed to further map cortico-striatal dysfunction in HD by using *in vivo* multimodal magnetic resonance imaging (MRI) techniques in the R6/1 HD mouse model. We took advantage of optogenetics tools (Zhang et al., 2010) and high-resolution cortico-striatal maps (Hintiryan et al., 2016) to modulate cortico-striatal function in HD mice. More specifically, we modulated the secondary motor cortex (M2) projection to dorsolateral striatum (DLS), whose structural alterations have been previously reported in HD (Hintiryan et al., 2016), and we used optogenetics coupled to *in vivo* microdialysis and to *ex vivo* multielectrode arrays (MEAs) to characterize cortico-striatal dysfunction in symptomatic HD mice. With this knowledge, we then aimed to test novel therapeutic interventions based on circuit restoration, an approach previously found successful for other basal ganglia disorders (Gradinaru et al., 2009; Kravitz et al., 2010).

Strikingly, selective optogenetic stimulation of M2-DLS afferent pathway successfully rectified motor learning and coordination deficits in symptomatic HD mice. All these effects were associated to improvements in synaptic plasticity such as an induced long-term depression (LTD) and a normalization of spine density within the striatum of HD mice. Our findings reveal that the function of M2 Cortex-DLS circuit is deeply impaired in the present HD mouse model and indicates that selective stimulation of this pathway induces long-lasting plasticity effects that significantly ameliorate motor symptoms in HD.

## Results

### Frontal cortex, but not motor cortex, show diminished functional connectivity with the striatum during rest in HD mice

To evaluate distinct cortico-striatal subcircuit alterations in the transgenic R6/1 mouse model of HD, we studied the functional connectivity using resting state fMRI in ∼20 week old R6/1 mice. First, the analysis of the whole–brain network showed reduced strength (p=0.0152) and global efficiency (p=0.0153) in HD mice compared to WT mice, as assessed by Student t test. In a more specific manner, we evaluated functional connectivity using seed-based analysis between frontal and motor cortices and selected basal ganglia regions (Figure 1); Supplementary files 1-4). In general, we found that frontal cortex and motor cortex from left hemisphere in HD mice showed positive correlation with a smaller brain area than the corresponding in WT mice. Then, we measured the mean correlation value for the regions of interest automatically identified based on a mouse brain atlas and genotype differences were analyzed (Figure 1b). Two-way ANOVA showed a significant brain region effect (F_(5,110)_ = 28.49 p<0.001), genotype effect (F_(1,22)_ = 11.96 p = 0.0022) and region/interaction effect (F_(5,110)_ = 3.064 p= 0.0125). In particular, compared to WT, the left frontal cortex of HD mice showed a significantly reduced correlation with left motor cortex (p = 0.0188), sensory-motor cortex (p = 0.0071), cingulate cortex (p = 0.0003), striatum (p = 0.0447), and with the thalamus (p= 0.0078), as shown by Bonferroni post hoc test. Conversely, mean correlation values between left motor cortex and the same basal ganglia nuclei were less affected. Indeed, two-way ANOVA showed a significant brain region effect (F_(5,110)_ = 51.99 p<0.0001), genotype effect (F_(1,22)_ = 9.278 p=0.00059) but not region/interaction effect (F_(5,110)_ = 2.196 p=0.0597). Bonferroni post hoc comparisons show significant differences with left frontal (p= 0.0176), sensory-motor (p= 0.0187) and cingulate cortices (p= 0.0041) and the thalamus (p= 0.0315). Thus, our results show that different cortico-striatal pathways are distinctly altered in HD mice, and particularly, fronto-striatal pathways, which include pre-motor areas in the mice, show stronger alterations than motor-striatal pathways at this symptomatic stage.

**Figure 1.**
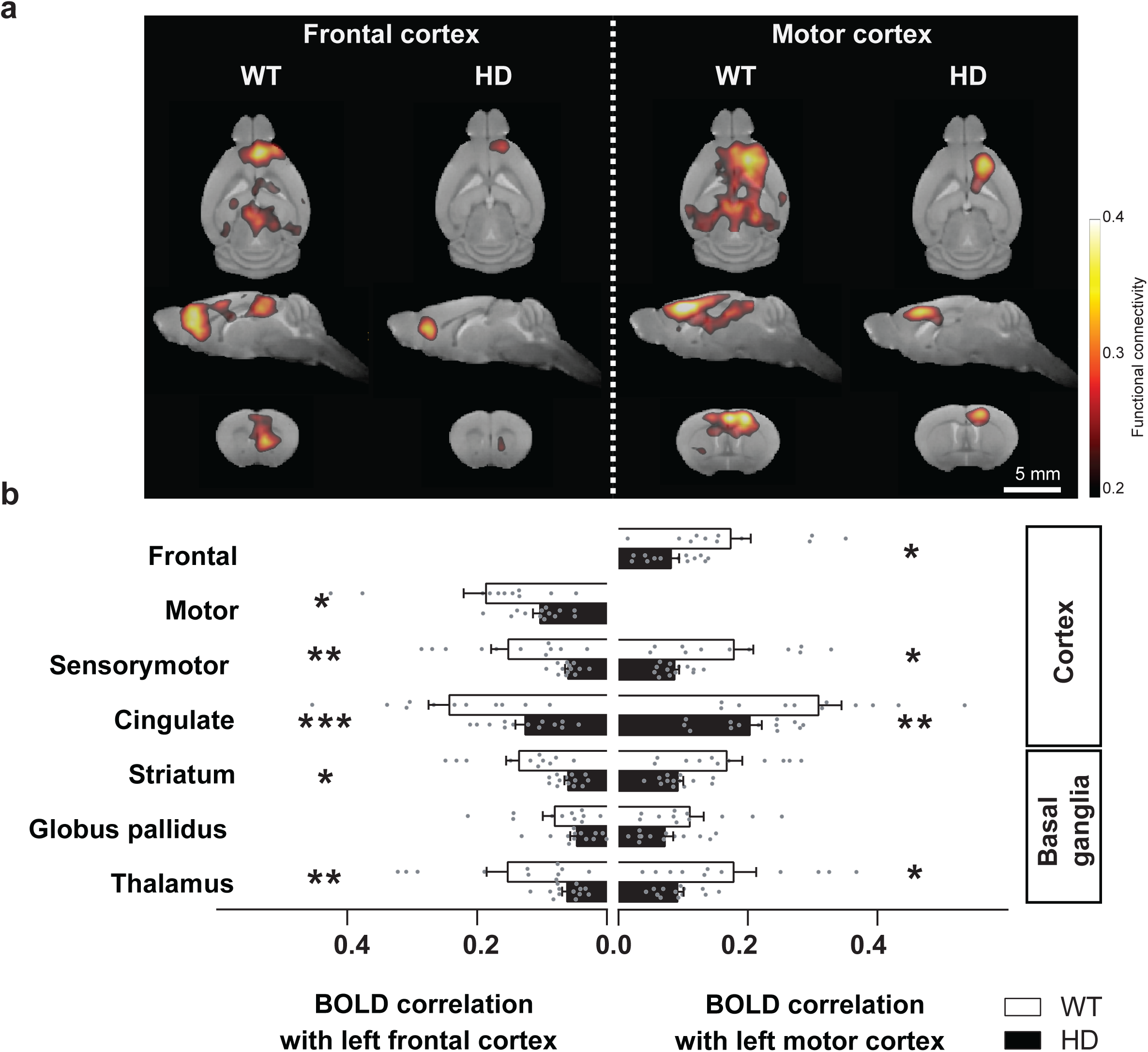
Frontal cortex and motor cortex functional connectivity is reduced in symptomatic HD mice. (**a**) Average seed-based BOLD correlation maps from frontal cortex (left panel) and motor cortex (right panel) in WT and the R6/1 mouse model of HD. (**b**) We measured the mean correlation value for cortical and basal ganglia in regions of interest obtained by atlas-based automatic parcellation. Both frontal cortex and motor cortex from HD mice showed positive correlation with smaller brain area than WT mice. Correlation between frontal and motor cortex and basal ganglia related structures from the left hemisphere are represented. Each grey point represents data from an individual mouse. Two-way ANOVA with Bonferroni post hoc comparisons test was performed. Data is represented as mean ± SEM (WT n=11 and R6/1 n=13 mice). *p<0.05, **p<0.01, ***p<0.001 HD *vs* WT.

We next assessed whether the functional differences found between these two cortical regions and the striatum were associated with changes at the microstructure level. Diffusion weighted imaging acquisition allowed the estimation of the fiber tracts connecting frontal cortex and striatum and motor cortex and striatum. Subsequent quantification of these tracts using diffusion metrics (fractional anisotropy) and tract-specific metrics (fiber-density) (Figure S1) showed that neither fractional anisotropy, nor fiber density were altered in these selected cortico-striatal circuits, indicating that at this stage mice show functional but not structural cortico-striatal deficits.

### Glutamate levels are reduced in the striatum of symptomatic HD mice

To further characterize cortico-striatal function in HD, we measured brain metabolites in the striatum of ∼17 weeks old mice, using single voxel ^1^H Magnetic Resonance Spectroscopy (MRS) (Figure 2). Two-way ANOVA showed metabolite-genotype interaction effect (F_(7,154)_ = 17.49 p<0.0001) and Bonferroni post hoc analysis of the absolute metabolites concentrations revealed similar glutamate and glutamine, increased creatine+phosphocreatine (Cr+PCr) (p = 0.0013), and decreased - N-Acethyl-aspartate (NAA) (p = 0.0371), NAA + NAA-Glutamate (NAAG) (p = 0.0057) and Taurine (p < 0.0001) (Figure 2b), in line with those previously described for the knock-in (KI) model of HD (Pépin et al., 2016). Moreover, the glutamate/glutamine ratio, which is related to glutamate released as a neurotransmitter, was reduced in HD mice (p = 0.0072, Student t-test), suggesting a reduced cortico-striatal function in HD mice (Figure 2c).

**Figure 2.**
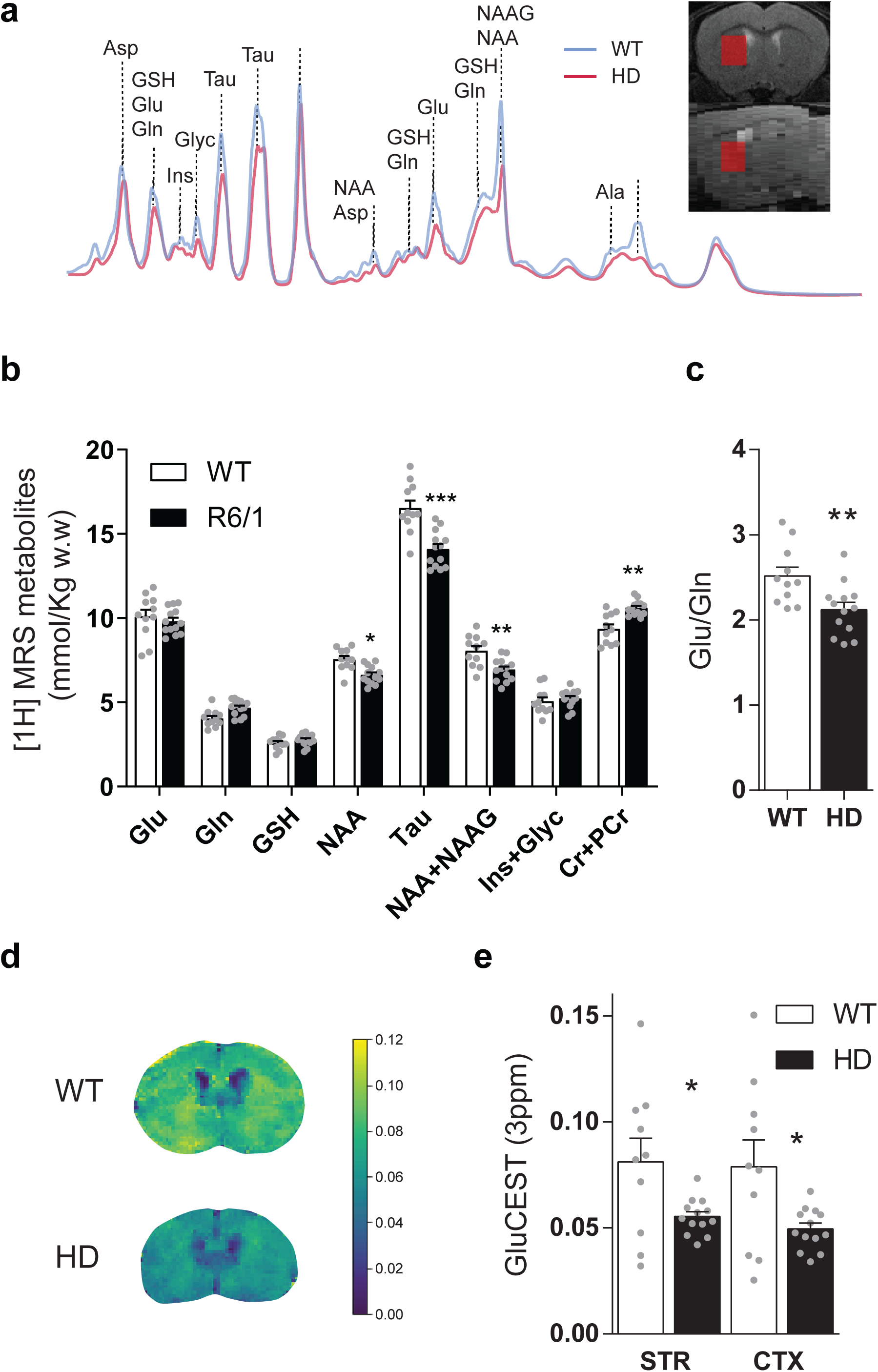
Brain metabolites in the striatum are altered in symptomatic HD mice. (**a**) Representative ^1^H MRS signal from abundant metabolites in the striatum of WT (blue line) and HD (red line) mice. Voxel location in the striatum is illustrated in coronal (top panel) and sagittal brain view (bottom panel). (**b**) Metabolites concentration quantification (mmol/Kg w.w). (**c**) Glutamate/Glutamine ratio was calculated as an indicator of glutamate neurotransmission (**d**) Representative coronal images of GluCEST in WT (top) and HD mice (bottom). (**e**) GluCEST quantification of striatum and cortex by manually drawn ROIs. Values were calculated as the GluCEST value (percentage of asymmetry at 3ppm) vs minimum GluCEST value in the ventricles. Data is represented as mean ± SEM (WT n=10 and HD1 n=13 mice). Each grey point represents data from an individual mouse. *p<0.05, **p<0.01, ***p<0.001 HD *vs* WT. Abbreviations: Glu: Glutamate; Gln: Glutamine; GSH: glutathione; NAA: N-Acethyl-aspartate; Tau: taurine; NAAG: N-Acethyl-aspartyl-glutamate; Ins+Glyc: Inositol + Glycine

In order to study the relationship between cortical and striatal levels of glutamate, we also used CEST (Chemical Exchange Saturation Transfer) technique, which has been demonstrated to map glutamate levels (Cai et al., 2013, 2012) with higher sensitivity and spatial resolution compared to ^1^H MRS (Bagga et al., 2018; Pépin et al., 2016). GluCEST showed decreased glutamate levels in striatum and cortex of HD mice compared to WT animals, as shown by two-way ANOVA brain region effect (F_(1,21)_ = 16.59 p = 0.0005), genotype effect (F_(1,21)_ = 6.541 p = 0.0183) but not region/interaction effect (F_(1,21)_ = 3.286 p = 0.0842), and Bonferroni post hoc comparisons showed significant differences between WT and HD in both regions (Figure 2d-e). Moreover, a significant correlation between MRS and GluCEST was observed for all cases (r= 0.5979; p=0.0026) (Figure S2).

### Selective M2 Cortex-striatal optogenetic stimulation reveals deficient glutamate release in HD mice

To functionally assess cortico-striatal function in HD mice, we evaluated optogenetically-induced glutamate release in the striatum of WT and HD mice using *in vivo* microdialysis (Figure 3a-c). After 6 baseline dialysate samples (collected every 6 min), blue light was ON for 5 minutes (during sample 7 collection) and 6 additional samples were collected in light-OFF conditions. Blue light stimulation led to increased glutamate levels in WT-ChR2 but neither in HD-ChR2 mice nor the WT-YFP and HD-YFP control mice. Glutamate levels returned to baseline after stimulation. Two-way ANOVA showed a significant effect of time (F_(12,144)_ = 2.219 p =0.0136) and Bonferroni post hoc analyses showed significant differences between WT-ChR2 and all other groups during stimulation (sample 7, blue box) (Figure 3c). These data indicate a reduced cortico-striatal-dependent glutamate release in symptomatic HD mice compared to WT controls. Glutamate concentration under baseline conditions was similar between genotypes (WT: 0.73 ± 0.08; R6/1: 0.69 ± 0.06 pmol/sample).

**Figure 3.**
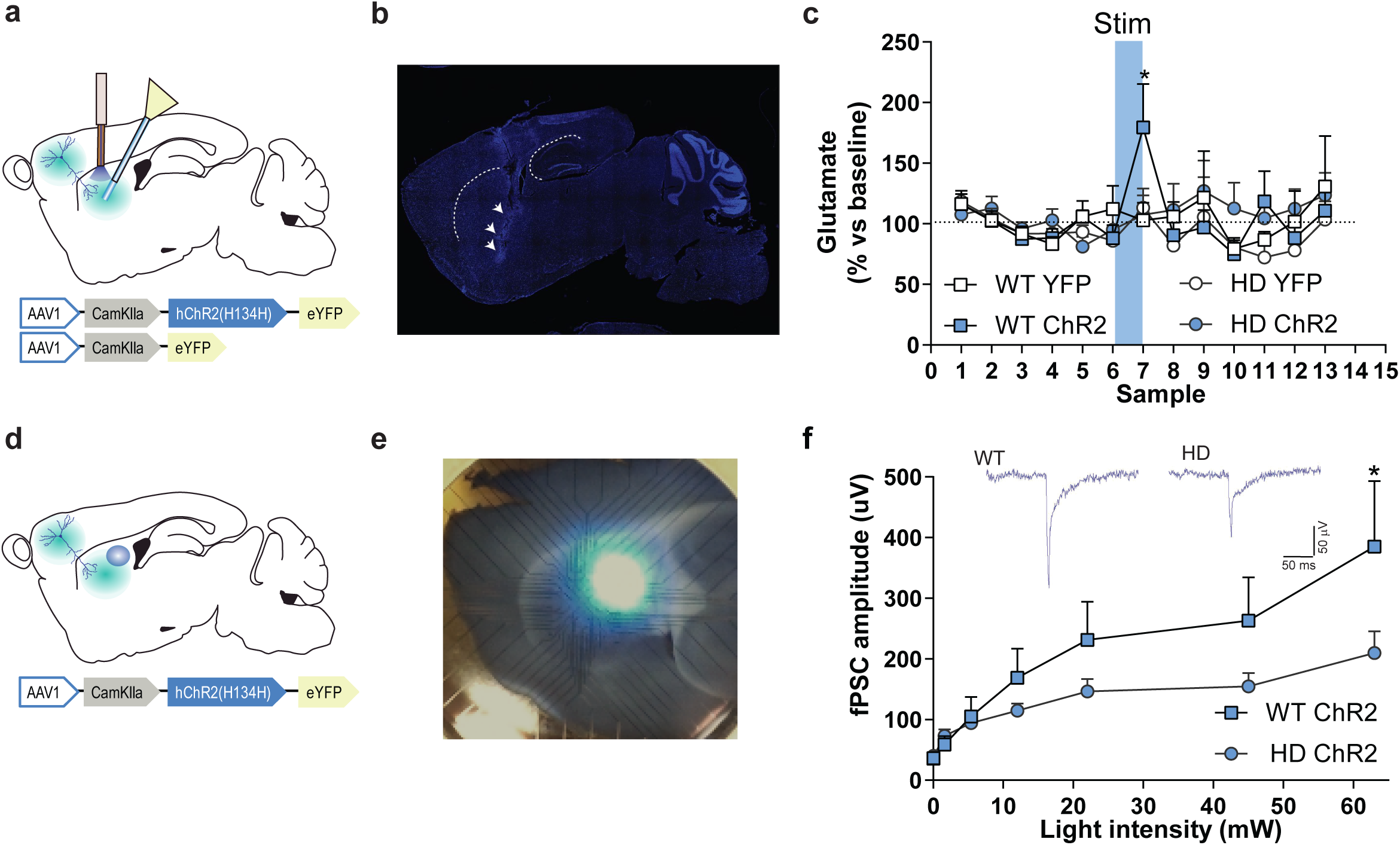
M2 Cortex-dorsolateral striatum function is impaired in HD mice. (**a**) Schematic representation showing AAV-ChR2 and control AVV-YFP constructs and injection location at M2 cortex and fiber-optic cannula implant and microdialysis probe in the dorsolateral striatum (DLS). (**b**) Representative histological image verification of microdialysis probe correct implant in the DLS. (**c**) Dialysate samples were collected every 6 min and glutamate was subsequently analyzed by HPLC. 473nm light stimulation was delivered at 10 Hz bilaterally for 5min in freely moving mice during sample 7 (blue box). Percentage of glutamate dialysate samples in the striatum respect mean baseline levels is represented as mean ± SEM (WT-YFP n=3, WT-ChR2 n=4, HD-YFP n=4, HD-ChR2 n=5 mice/group). (**d**) Schematic representation of multielectrode array recordings in slices. AAV-ChR2 was injected at M2 cortex. Sagittal slices were obtained for light induced recording. Fiber-optic cannula was used to locate the light on top of the dorsolateral striatum. (**e**) Representative image showing electrodes and location of light stimulation. (**f**) Amplitude of light intensity-induced striatal field postsynaptic currents (fPSC) in WT and HD mice. Inset: Representative rasters of the light-induced electrical response. Two-way ANOVA with Bonferroni post hoc comparisons test was performed. Values are expressed as mean ± S.E.M (WT-ChR2 n=4, HD-ChR2 n=3 mice). *p<0.05.

### Striatal neuronal response to M2 Cortex afference stimulation is dampened in HD mice

To test whether optical stimulation was effectively activating the cortico-striatal pathway, we performed multielectrode array (MEA) recordings in sagittal acute brain slices of WT-ChR2 and HD-ChR2 mice (Figure 3d-f). We recorded fPSC in the dorsal striatum after stimulation with increasing light intensities to create a normalized input-output assay. We found that optic stimulation of M2 Cortex afferent pathways evoked fPSC in the dorsal striatum (Figure 3f). Two-way ANOVA showed that the amplitude of the evoked striatal fPSC was significantly larger in WT-ChR2 than in HD-ChR2 mouse (genotype effect; F_(10,55)_ = 1.188; p = 0.0395) followed by Bonferroni post hoc comparisons. Also, increasing laser intensity led to fPSC of larger amplitude in both mouse groups (light intensity factor; F_(10,55)_ = 9.906; p < 0.00001. These results confirm the reliability of our optogenetic approach since they provide evidences of a specific striatal response to the selective optogenetic activation of cortico-striatal afferences. In addition, the results further confirm that M2 Cortex-striatal circuit in HD is altered, possibly due to both pre and post synaptic effects.

### Repeated M2 Cortex-DLS stimulation improves exploratory and stereotypic behavior in symptomatic HD mice

We next examined whether the increased glutamatergic input onto DLS -induced by optogenetic stimulation of M2 afferences-translated into an improved motor behavior in HD mice. To this end, we performed repeated optogenetic cortico-striatal stimulation in freely moving mice in the open field (OF) apparatus (Figure 4), similar to Ahmari et al 2013. Mice were placed in the OF and, after 5 min, blue light was delivered during 1 min at 10 Hz, and mice left in the apparatus for 5 extra minutes. Distance travelled, rearing time and stereotypic behavior were recorded during this 11 min period (Figure 4b, 4d). The same procedure was repeated 3 times, each one-week apart, in symptomatic 20, 21 and 22-week old mice. The precision of the injection was evaluated by YFP expression at M2 Cortex injection site and the presence of fluorescent fibers in the striatal region (Figure 4c; Figure S3).

**Figure 4.**
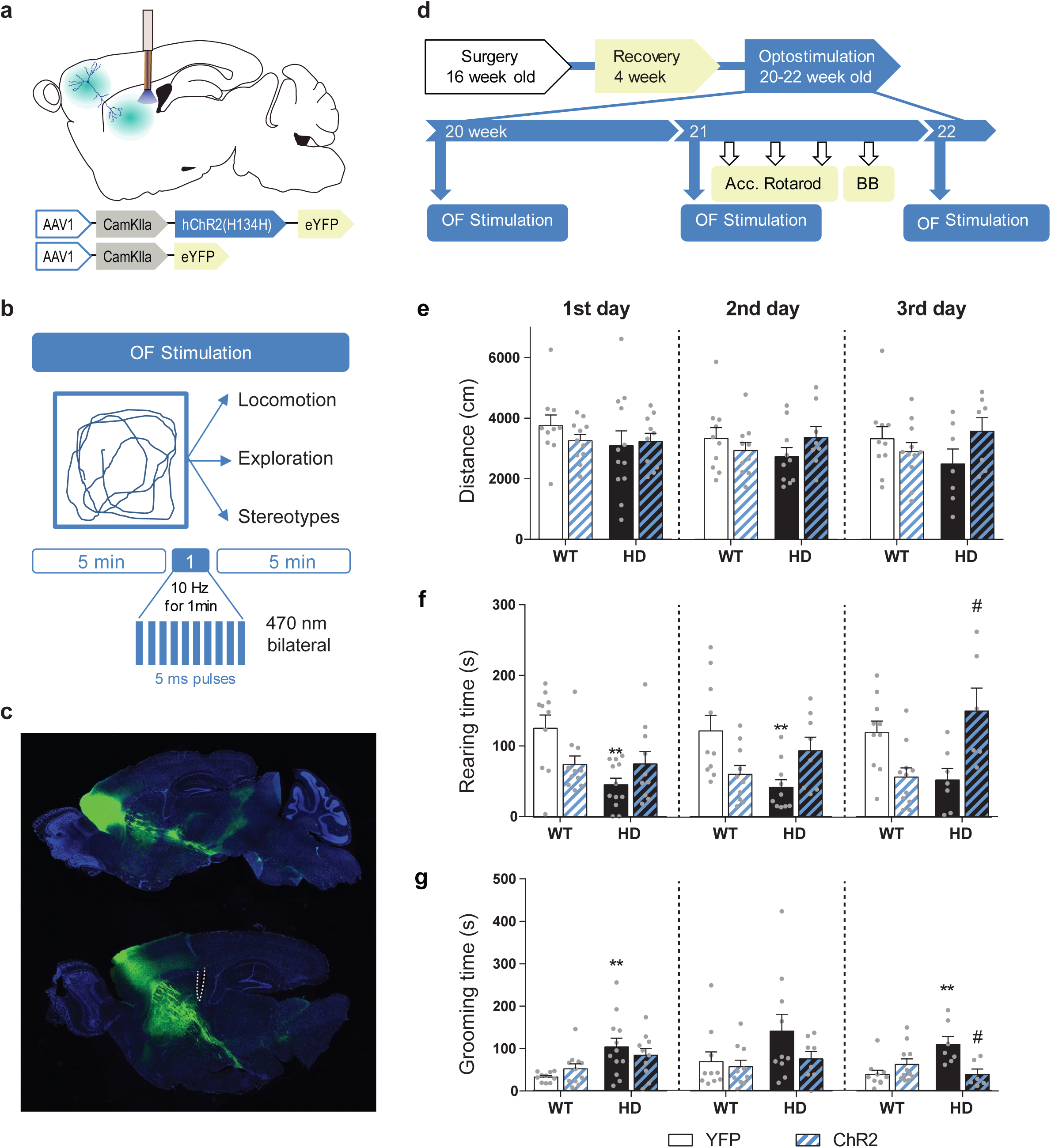
Repeated optogenetic stimulation of the cortico-striatal pathway recover exploratory and stereotypic behavioral deficits in symptomatic HD mice. (**a**) Schematic diagram showing AAV-ChR2 and control AAV-YFP constructs injections at M2 cortex and fiber-optic cannula implant in the DLS allowing light stimulation of cortically infected neuronal axons. (**b**) Locomotion (distance travelled), exploration time (rearing time) and stereotypes (grooming time) were evaluated during 11 min open field session Optogenetic stimulation consisted in 10 Hz light stimulation for 1 min during open field (OF) task. (**c**) Representative fluorescence image showing the precision of the AAV injection by YFP expression at M2 Cortex and the presence of YFP fibers in the striatal region. (**d**) Surgery, behavior and optogenetic experimental timeline. The OF procedure was performed at 20 (1^st^ day OF), 21 (2^nd^ day OF) and 22 (3^rd^ day OF) week old mice. After 2^nd^ stimulation day, motor learning and coordination tests were performed. (**e**) Distance travelled, (**f**) rearing time and (**g**) stereotypic behavior were measured during the 11 min OF period. Values are expressed as mean ± S.E.M (WT-YFP n=10, WT-ChR2 n=11, HD-YFP n=13 and HD-ChR2 n=11). Each point represents data from an individual mouse. Data were analyzed by two-way ANOVA with genotype and light stimulation as factors, and Bonferroni test as a post hoc. *p<0.05, **p<0.01 *vs* WT-YFP and #p<0.05 *vs* HD-YFP

Total distance travelled was similar in all experimental groups during OF sessions (Figure 4e) and light stimulation did not induce significant changes in locomotor activity during the 1 min stimulation (Figure S4).

In contrast, exploratory behavior, measured as rearing time, was reduced in HD mice compared to WT, as previously described for KI mice (Pépin et al., 2016) and optogenetic stimulation induced opposite change in WT-ChR2 and HD-ChR2 mice, with an increase of exploratory behavior over the sessions in HD-ChR2 mice (Figure 4f). Two-way ANOVA with genotype and stimulation as factors showed significant differences between groups (First day: genotype F _(1, 39)_ = 7.540 p =0.0091 and interaction effect F _(1, 39)_ = 7.889 p = 0.0077; Second day: Interaction effect F _(1, 34)_ = 11.56 p = 0.0017; Third day: Interaction effect F _(1, 30)_ = 17.90 p<0.0004. Bonferroni post hoc analyses were performed and revealed significant differences between WT-YFP and HD-YFP during first and second OF sessions and between HD-YFP and HD-ChR2 during the third OF session. Total number of rearing events showed similar results (Figure S4b).

In addition, we measured stereotypic behavior, mainly grooming, in the same mice (Figure 4g, S4c). Grooming is a maintenance behavior suggestive of mice wealth, but also associated with stereotypic pathological behavior when is substantially increased. In non-stimulated conditions, HD mice showed an increased stereotypic behavior, as compared to WT mice, in line with the previously described increase in grooming in KI mice (Pépin et al., 2016). Interestingly, repeated optogenetic stimulation reduced stereotyped behavior in HD-ChR2 mice. Two-way ANOVA revealed significant differences between groups (First day: genotype effect F _(1, 39)_ = 11.36 p= 0.0017; Third day: Interaction effect F _(1, 30)_ = 11.42 p =0.020). Bonferroni post hoc analyses revealed significant differences between WT-YFP and HD-YFP during first and third OF session and between HD-YFP and HD-ChR2 during the last OF session. Total number of grooming events showed similar results (Figure S4c).

Our data reveal that M2 Cortex-striatal stimulation has a stronger impact in the modulation of spontaneous activity of HD mice while acute locomotor activation was not observed. The effects of the light stimulation on exploratory behavior are manifested before any stimulation during the 2^nd^ and 3^rd^ day session, suggesting there are some long-lasting effects evoked by the optogenetic stimulation of this M2-DLS circuit.

### Motor learning and coordination deficits in symptomatic HD mice are recovered by optogenetic stimulation of the M2-DLS cortico-striatal pathway

We further explored the effects of M2-DLS stimulation on motor learning and coordination. Motor learning was assessed by measuring the latency to fall from an accelerating rotarod. HD mice showed reduced latency to fall compared to WT, as previously described (Creus-Muncunill et al., 2019; Puigdellívol et al., 2015). Interestingly, optogenetic stimulation of the cortico-striatal pathway was able to increase the latency to fall from the rod in HD-ChR2 mice to similar levels than WT mice. Repeated measures ANOVA showed significant effects of mice group F _(3, 35)_ = 6.058; p<0.0020, time F _(11, 385)_ = 24.24, p<0.0001 and interaction F _(33, 385)_ = 2.160; p<0.0003. Moreover, Bonferroni post hoc analyses showed significant differences between WT-YFP and HD-YFP, which are mostly lost in HD-ChR2 (Figure 5c). In addition, motor balance and coordination were evaluated by measuring the ability of mice to walk across a balance beam during 2 min (Figure 5d). The number of 5 cm-frames crossed was reduced in HD-YFP mice compared to WT mice, as already described (Creus-Muncunill et al., 2019). Remarkably, cortico-striatal stimulation ameliorated motor coordination in HD-ChR2 mice, reaching similar levels to WT mice. Two-way ANOVA showed a genotype effect (F_(1, 34)_ = 7.226; p<0.011), with no stimulation (F_(1, 34)_ = 1.270; p = 0.267) or interaction effect F_(1, 34)_ = 2.863; p = 0.0998, and Bonferroni post hoc analyses showed significant differences only between WT-YFP and HD-YFP mice groups. These data suggest that the prior activation of M2-DLS circuit induces long-lasting effects that impact in the basal ganglia motor function. Moreover, our results indicate that by restoring selected cortico-striatal circuits, i.e. M2-DLS circuit, we can successfully ameliorate motor function in HD.

**Figure 5.**
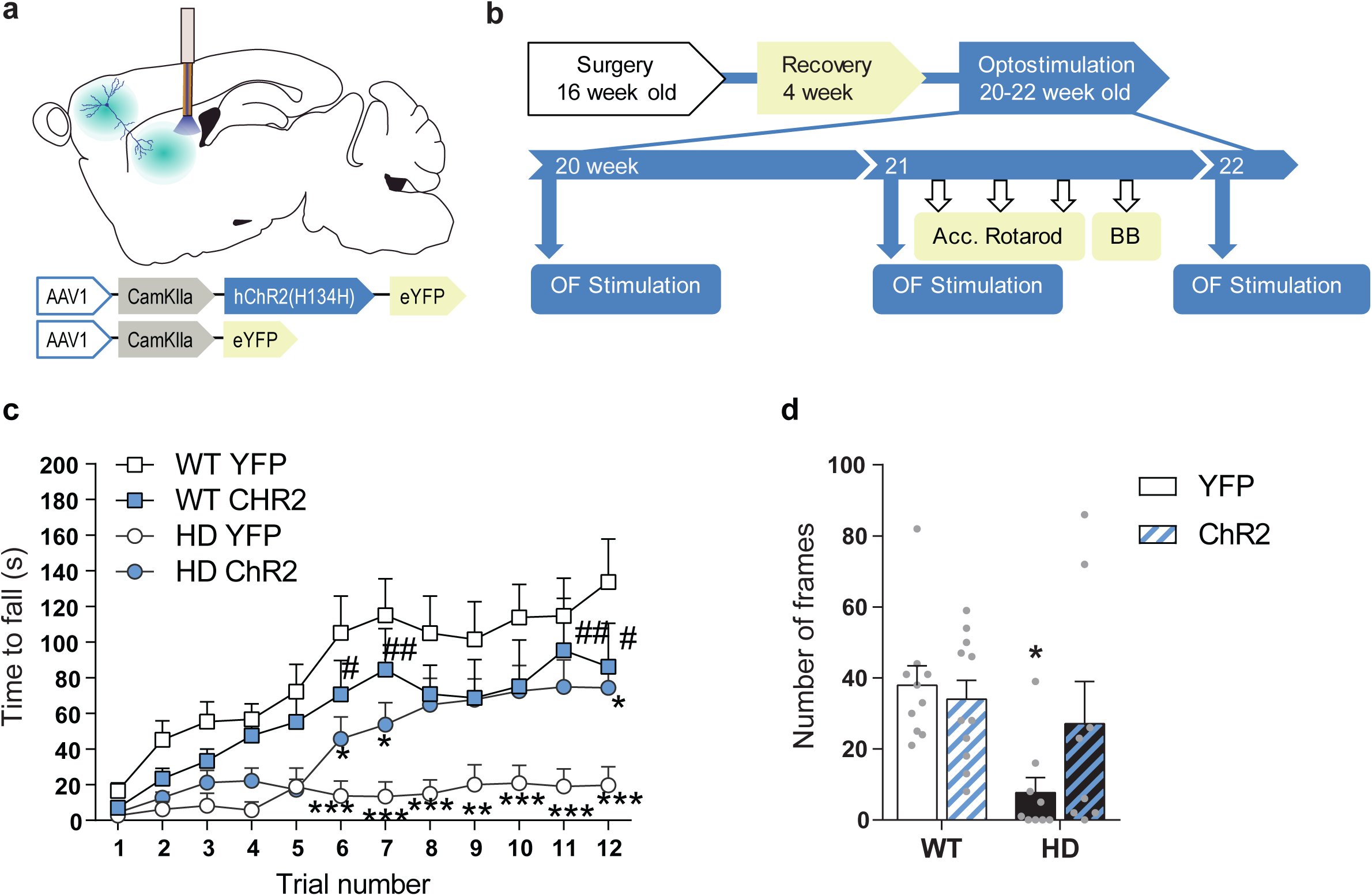
Optogenetic stimulation of the M2 Cortex-DLS pathway recover motor learning and coordination deficits in HD mice. (**a**) Schematic representation showing AAV-ChR2 and control AVV-YFP constructs injections at M2 cortex, fiber-optic cannula implant in the DLS. (**b**) Surgery, behavior and optogenetic experimental timeline. After 2^nd^ stimulation day in the OF, motor learning (accelerating rotarod) and coordination (balance beam, BB) tests were performed. (**c**) Latency to fall in the accelerating rotarod task. (**d**) Number of frames crossed in the BB test. Each point represents data from an individual mouse. Data were analyzed by repeated measures ANOVA with group and time as factors for accelerating rotarod and by two-way ANOVA with genotype and stimulation as factors for BB test. Bonferroni’s Multiple Comparison test was performed as post hoc test *p<0.05, **p<0.01, ***p<0.001 vs WT-YFP and #p<0.05 vs HD-YFP. Values are expressed as mean ± S.E.M (WT-YFP n=10, WT-ChR2 n=11, HD-YFP n=10 and HD-ChR2 n=9).

### Repeated cortico-striatal stimulation triggers persistent improvements of synaptic plasticity in symptomatic HD mice

We explored whether these behavioral effects where due to long-lasting synaptic re-wiring of cortico-striatal circuit in HD (Figure 6). We repeated the cortico-striatal stimulation procedure in vivo (Figure S5) and then explored if the repeated optogenetic stimulation modulates the electrophysiological response in cortico-striatal slices. First, we confirmed that MSNs responded to light stimulation. As previously done, we recorded fPSC response to increasing light intensity stimulation. We found that increasing laser intensity led to striatal fPSC of increasing amplitude in both WT-ChR2 and HD-ChR2 groups as shown by two-way ANOVA light intensity effect (F_(7,154)_ = 39.05; p<0.0001). Interestingly, we found no genotype effect or genotype/light intensity interaction (Figure 6b).

**Figure 6.**
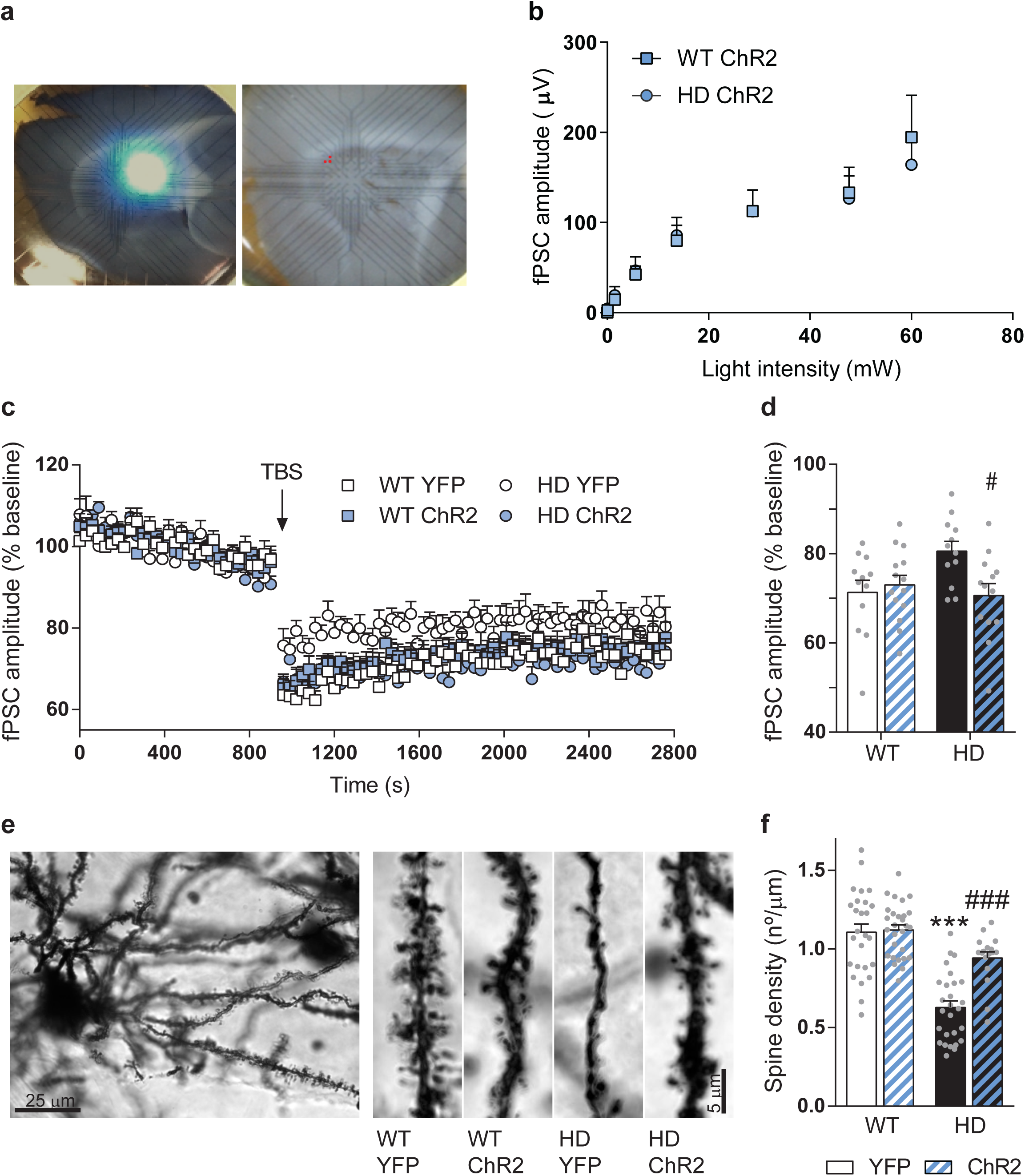
Repeated optogenetic cortico-striatal stimulation restores synaptic plasticity and dendritic spine density in HD mice. (**a**) Representative image were location of the light stimulation (left image) and stimulating electrodes (right image, highlighted in red) are shown. (**b**) Light intensity-induced striatal field postsynaptic currents in WT and HD mice with AAV-ChR2 expressed in M2 Cortex. (**c**) The graph shows the time course of LTD evoked at cortico-striatal synapses following a TBS. The TBS, indicated by the arrow, was presented after 15 min of baseline recordings. Field postsynaptic currents (fPSC) are represented as a percentage of baseline. (**d**) Histogram shows the averaged amplitude of fPSC evoked during 30 min after TBS. Data were analyzed by repeated measures ANOVA with genotype and stimulation as factors and Bonferroni’s Multiple Comparison as post hoc test *p<0.05, **p<0.01, ***p<0.001 vs WT-YFP and #p<0.05 vs HD-YFP. Values are expressed as mean ± S.E.M (WT-YFP n=12, WT-ChR2 n=13, HD-YFP n=12 and HD-ChR2 n=14 mice). (**e**) Golgi-impregnated representative neuron and segments of secondary dendrites from MSNs from WT and HD mice infected with AAV-YFP and AAV-ChR2 respectively. Scale bar, 3 μm. Dendritic spines were counted in a segment of known length (∼20μm) to obtain the spine density. (**f**) Quantitative analysis of spine densities per micrometer of dendritic length of 3-13 neurons per mice. HD-YFP mice exhibit a significant reduction in dendritic spines that was significantly increased in stimulated HD-ChR2. Two-way ANOVA with Bonferroni’s post hoc comparisons test was performed. All data are shown as the mean ± S.E.M.

Then, we recorded fPSC in the striatum after electrical stimulation of cortical afferences. We induced LTD, which is known to be altered in HD mouse (Creus-Muncunill et al., 2019), using a theta-burst stimulation (TBS) protocol on cortical afferences and compared the amplitude of evoked striatal fPSC (Figure 6c-d). Both, WT-YFP and WT-ChR2 mice, presented a significant decrease in the amplitude of evoked fPSC following the LTD protocol that lasted at least for 30 min, indicating cortico-striatal LTD induction. In the same conditions, stable LTD was induced in HD-ChR2 animals, while only a slight LTD was induced in HD-YFP. Two-way ANOVA showed a mice group/time interaction (F_(273,4277)_ = 1.573; p < 0.0001), time effect (F_(91,4277)_ = 111.9; p < 0.0001) and group effect (F_(3,47)_ = 3.312; p = 0.0279) and main group effects showed differences between HD-YFP and HD-ChR2 (p=0.04), as shown by Bonferroni post hoc test. Indeed, similar results are obtained when analyzing the average of the fPSC amplitude after TBS (Figure 6d). Here, two-way ANOVA showed a genotype/stimulation interaction (F_(1,47)_ =5.621; p = 0.0219) and Bonferroni post hoc test showed that optogenetic stimulation restored the ability to induce LTD in HD mice.

To examine whether improved cortico-striatal synaptic function correlates with plasticity changes, we evaluated spine density from Golgi-impregnated striatal MSNs (Figure 6e-f). Spine density in striatal MSN from HD mice was reduced compared to WT mice, as previously shown for 3-month old YAC128 (Marco et al., 2013) and in hippocampal apical dendrites from R6/1 (Miguez et al., 2015). Repeated optogenetic stimulation restored spine density in HD mice, without altering spine density in WT mice, as indicated by 2-way ANOVA with genotype F _(1, 94)_ = 56.52 p<0.001; stimulation F _(1, 94)_ = 14.29 p<0.0003 and interaction effect F _(1, 94)_ = 11.77 p<0.0009. Bonferroni post hoc comparisons showed significant differences between WT-YFP and HD-YFP (p < 0.0001), and between HD-YFP and HD-ChR2 (p < 0.0001), indicating that repeated optogenetic stimulation of the cortico-striatal pathway restores spine density in the striatum of HD mice.

Altogether, our results show that optogenetic M2-striatal stimulation induces persistent synaptic plasticity changes that could explain the behavioral improvements found in HD mice.

## Discussion

Basal ganglia circuit dysfunction is the main cause of motor abnormalities in HD. In particular, cortico-striatal disconnection has a prominent role in the disease progression, however, specific cortico-striatal subcircuits have not been explored in detail in relation to HD. Here, we further characterized cortico-striatal functional alterations in the symptomatic HD mouse. Our data reveal that particularly M2-DLS pathway is profoundly compromised in our R6/1 mouse model of HD. The idea to treat brain circuits using optogenetic tools is object to study in many neurological disorders, such as Parkinson (Gradinaru et al., 2009; Kravitz et al., 2010; Magno et al., 2019), stroke (Cheng et al., 2014; Tennant et al., 2017) and psychiatric disorders (Fuchikami et al., 2015). In this line, we aimed to restore circuit function in HD by increasing selected cortico-striatal activity in symptomatic mouse. Indeed, we provide strong evidence that we can ameliorate HD symptoms in symptomatic HD mice by stimulating selected M2 cortex-DLS pathway.

### Fronto-striatal circuits show broader functional alterations than motor-striatal circuits in HD

Cortico-striatal function is known to be impaired along the progression of HD. However, little is known about how specific cortico-striatal subcircuits are affected. Here, we showed that fronto-striatal functional connectivity is impaired in our HD symptomatic mice, while motor-striatal connectivity is spared. Our results indicate that cortical afferences rather than output pathways are predominantly affected during disease progression, highlighting that frontal cortex, rather than primary motor cortex, has a stronger contribution to the basal ganglia network dysfunction in HD. These results in symptomatic mice are in line with previous data in prodromal HD patients, where correlation between caudate nucleus with premotor areas, but not primary motor cortex, was affected (Unschuld et al., 2012). Indeed, topographically selective changes in the cortex explains the clinical heterogeneity found in HD. For example, larger primary motor cortex degeneration correlates with motor impairments while cingulate cortex degeneration correlates with predominant mood symptomatology (Estrada-Sánchez and Rebec, 2013; Rosas et al., 2008; Thu et al., 2010). Therefore, it is relevant to distinguish between different subcortical regions when evaluating HD disease progression and symptoms, and more importantly for therapeutic interventions.

### M2-DLS glutamate transmission is reduced in symptomatic HD mice

There is a controversy in the literature regarding whether total levels of glutamate are altered in HD. Cortical and striatal metabolites, measured by ^1^H MRS, are known to be profoundly altered in symptomatic HD models (Zacharoff et al., 2012), and glutamate concentration, measured by GluCEST, is known to be reduced in different brain regions, including most of the basal ganglia nuclei involved in motor behavior (Pépin et al., 2016). In our R6/1 mouse model, brain metabolites, measured using ^1^H MRS, showed significant changes in line to the ones observed for KI (Pépin et al., 2016), R6/2 (Tkac et al., 2007) and *in vitro* MRS striatal extracts from R6/2 mice (Jenkins et al., 2000). However, when evaluating absolute values by ^1^H MRS, we failed to show total glutamate alterations in the striatum of HD mice. Nevertheless, unaltered glutamate levels were also described in R6/2 mice (Zacharoff et al., 2012). Still, glutamate/glutamine levels, which ratio is linked to functional/released glutamate, is reduced in the striatum of HD mice, indicating that glutamate transmission is defective in HD.

We then evaluated glutamate release from cortical terminals, as cortical afferences provide the major glutamatergic source to the striatum. Optogenetic stimulation of cortical afferences in the striatum induced detectable glutamate increase in WT mice, but not HD mice, clearly indicating deficits in glutamate release in symptomatic stages. This effect is further supported by the blunted electrophysiological response to M2 optogenetic acute stimulation observed in the striatum of HD mice. Although previous work described higher glutamate release in R6/1 mice (NicNiocaill et al., 2001), these discrepancies could be explained by the different location of the microdialysis probe (dorsomedial striatum (DMS) *vs* DLS), mice age (16 *vs* 20 weeks old mice), and strategy to induce glutamate release (selective optogenetic stimulation *vs* KCl or NMDA application), which makes difficult to evaluate and compare sources of the glutamate measured (cortico-striatal *vs* thalamic or even astrocytic).

Moreover, convergent evidence indicates impaired glutamate release in HD, although most of them rely in electrophysiological measurements (Estrada-Sánchez and Rebec, 2013; Klapstein et al., 2001; Milnerwood and Raymond, 2007). Indeed, the monitoring of synaptic changes during the progression of HD in the YAC128 model reported biphasic age-dependent alterations of the cortico-striatal activity (Joshi et al. 2009). Asymptomatic HD mice present increased glutamate release, postsynaptic currents and synaptic responses. However, there is a shift in the three parameters as symptoms appear which are maintained at late stages. Thus, symptomatic YAC128 mice exhibit reduced glutamate release, post synaptic currents and synaptic responses. All together, these data indicate that reduced glutamate release from the cortex, at least in late stages of the disease, might lead the basal ganglia network alterations found in HD patients.

### Motor alterations in HD involve specifically M2-DLS cortico-striatal dysfunction

Motor learning and coordination has been extensively characterized in animal models of HD (Creus-Muncunill et al., 2019; Hong et al., 2012; Puigdellívol et al., 2015). In this work, we particularly support the idea that M2-DLS activity is necessary for the proper execution of motor learning tasks, as by manipulating this specific cortico-striatal subcircuit we restore motor defects in pathological HD mice. Indeed, primary and secondary motor cortex have different contributions to action differentiation and sequencing in mice. M2 lesions impairs acquisition and reversal of a simple sequence consisting in two lever presses, one distal and the other proximal to reward (Yin, 2009), while M1 lesions only impairs reversal. Thus, M2 activity seems more related to motor learning than M1. In fact, there is an extensive functional reorganization in the striatum during different phases of learning during a motor learning task, such as accelerated rotarod (Yin, 2009) or serial order task (Rothwell et al., 2015). This includes initial predominant activity in DMS during early training, while extensive training engages DLS, indicating that DLS has a crucial role in completion of a learned sequence (Rothwell et al., 2015; Yin, 2009). Moreover, activity of MSNs from direct or indirect output pathways in the DLS also experience divergent firing patterns during a motor learning task (Jin et al., 2014). Action initiation correlates to increased activity in both dMSNs and iMSNs in DLS, and, as the action sequence progress, most iMSNs reduce firing rate while dMSNs exhibit sustained activity (Jin et al., 2014; Rothwell et al., 2015). Taking this together, we suggest that our M2-DLS optogenetic stimulation might strengthen the ability to complete a motor learning task in HD mice.

In this line, the proportion of cortico-striatal inputs to the direct or indirect pathways depends on the cortical subregion from where these inputs arise (Wall et al., 2013). For example, primary motor cortex send axonal projections in greater extend to the indirect pathway, while sensory cortex projects more to direct pathway and secondary motor cortex project similarly to both striatal output circuits (Wall et al., 2013). In addition, in the cortex there are also sparse long-range GABAergic projection neurons that modulate striatal output and motor behavior (Melzer et al., 2017; Rock et al., 2016). Specifically somatostatin, but not PV, GABAergic neurons from M2 inhibit movement by both D1 and D2 MSNs inhibition, while somatostatin GABA long-range projections from M1 increase movement by inhibiting cholinergic interneurons in the striatum (Melzer et al., 2017). Thus, there is an urgent need to precisely map region and cell-specific striatal inputs to understand the functional organization of the cortico-striatal subcircuits and its impact on behavior, both under physiological and pathological conditions.

### Repeated M2 Cortex-DLS stimulation lead to sustained synaptic plasticity changes and rectify motor deficits in HD

Our data demonstrate that selective optogenetic stimulations induce long-lasting effects in neuronal plasticity and behavior, which are fundamental for brain re-wiring in pathological conditions. Optogenetic stimulation of infralimbic prefrontal cortex induces sustained antidepressant behavioral and synaptic effects (i.e. spine density) (Fuchikami et al., 2015), and optogenetic stimulation of motor cortex promotes functional recovery after stroke (Cheng et al., 2014). Also, repeated cortico-striatal optogenetic stimulation generates persistent stereotypic behavior in mice (Ahmari et al., 2013) that lasts at least 2 weeks, which is specifically dependent of the cortico-striatal subcircuit stimulated (i.e. orbitofrontal cortex (OFC)–ventromedial striatum (VMS)) (Ahmari et al., 2013). In contrast to our results, OFC– VMS stimulation induces grooming behavior in mice (Ahmari et al., 2013), pointing that the specific effects are dependent of the distinct cortico-striatal subcircuits.

More surprising was the finding that M2-DLS stimulation was able to ameliorate exploratory behavior and stereotypes in HD mice. Our data show a decrease in rearing and an increase in grooming time in our R6/1 mice compared to WT, as was previously observed in KI mice (Hong et al., 2012; Pépin et al., 2016). In addition, we were able to induce exploratory activity, measured as rearing time, and reduce stereotypic behavior, mainly grooming, in already symptomatic mice by M2-DLS optogenetic stimulation. Active exploration and grooming modulate cortico-striatal phase synchrony in R6/2 mice (Hong et al., 2012), further indicating that cortico-striatal function is involved in these behaviors. In here, we further demonstrate that these spontaneous behaviors can be modulated by M2-DLS pathway, and again, the effects are persistent at least one weak, as changes are already observed prior the light stimulation in the second and third stimulation day. Therefore, optogenetic manipulations at cortical circuits generate sustained synaptic plasticity changes. However, if the long-lasting effects are specific of cortical circuits or is a general feature of optogenetic stimulation remains to be determined.

Overall, we identified fronto-striatal circuits, and particularly the M2-DLS cortico-striatal subcircuit to be profoundly altered in symptomatic HD mice. Moreover, by taking advantage of optogenetic techniques, we demonstrated altered glutamate release from M2 cortical projection neurons and deficient response in the striatum in HD. In future studies, the ability to dissect the function of different cortico-striatal subcircuits in physiological and pathological conditions might help to further understand how mHtt alters the information processing in the basal ganglia. Here, we highlight a key role of the M2-DLS specific subcircuit in the modulation of motor alterations in HD, including exploratory behavior, stereotypes, motor learning and coordination. In this context, we were able to successfully reestablish physiological, morphological and behavior alterations in our symptomatic HD mice by generating persistent synaptic plasticity in selected cortico-striatal circuits. Our successful approach opens new opportunities to design therapeutic strategies to ameliorate HD symptoms based on circuit restoration, which might be useful also for other basal ganglia disorders.

## Materials and Methods

### Animals

R6/1 transgenic mice expressing exon-1 of mutant huntingtin containing 115 CAG repeats were acquired from Jackson Laboratory (Bar Harbor, ME, USA) and maintained in a B6CBA background. Genotypes were obtained by polymerase chain reaction (PCR) from tail biopsy and WT littermates were used as control group. Animals were housed together in groups of mixed genotypes in a room kept at 19-22°C and 40-60% humidity under a 12:12 h light/dark cycle with access to water and food ad libitum, and data were recorded for analysis by microchip mouse number. All animal procedures were approved by the animal experimentation Ethics Committee of the Universitat de Barcelona (274/18) and Generalitat de Catalunya (10101), in compliance with the Spanish RD 53/2013 and European 2010/63/UE regulations for the care and use of laboratory animals.

### MRI Image acquisition

11 WT mice and 13 R6/1 mice were scanned at 17-20 weeks of age on a 7.0T BioSpec 70/30 horizontal animal scanner (Bruker BioSpin, Ettlingen, Germany), equipped with an actively shielded gradient system (400 mT/m, 12 cm inner diameter). Each animal underwent two acquisition sessions. First session included magnetic resonance spectroscopy (MRS) and CEST (chemical exchange saturation transfer) acquisition to assess metabolite concentration. The second session was performed one week later and included structural T2-weighted imaging, diffusion weighted imaging (DWI) and resting-state functional magnetic resonance (rs-fMRI) to evaluate connectivity between regions of interest. In both cases, animals were placed in supine position in a Plexiglas holder with a nose cone for administering anesthetic gases and were fixed using tooth an ear bars and adhesive tape. Animals were anesthetized with 2.5% isoflurane (70:30 N2O:O2). In the second session, a combination of medetomidine (bolus of 0.3 mg/kg, 0.6 mg/kg/h infusion) and isoflurane (0.5%) was used to sedate the animals.

In both sessions, 3D-localizer scans were used to ensure accurate position of the head at the isocenter of the magnet. During the first session, MRS was acquired using PRESS (voxel size 5.4 µl, TR=5000ms, TE=12ms, 256 averages, partial water suppression, VAPOR) in a voxel located in left striatum (Figure 2a). Voxel position was defined using T2 RARE images acquired in axial, sagittal and coronal orientation as reference. A non-suppressed reference water signal was also acquired in the same voxel (8 averages). GluCEST and WASSR (water saturation shift referencing) were acquired from one coronal slice with slice thickness 2.5 mm, covering striatum and cortex. WASSR image was acquired as reference using a pulse of 500 ms and 0.5 µT with an offset range from -200 and 200 Hz. GluCEST sequence used a pulse of 6000ms, 10µT and offset ranging from -1600 to 1600 Hz.

In the second session, T2-weigthed image was acquired using a RARE sequence with effective TE=33ms TR=2.3 s, RARE factor=8, voxel size=0.08×0.08 mm^2^ and slice thickness=0.5 mm. DWI was acquired using an EPI sequence with TR= 6 s, TE=27.6 ms, voxel size 0.21×0.21 mm^2^ and slice thickness 0.5mm, 30 gradient directions with b-value=1000 s/mm^2^ and 5 baseline images (b-value=0 s/mm^2^). rs-fMRI was acquired with an EPI sequence with TR=2s, TE=19.4, voxel size 0.21×0.21 mm^2^ and slice thickness 0.5mm. 420 volumes were acquired resulting in an acquisition time of 14 minutes.

### Structural connectivity analysis

The fiber tracts connecting 1) frontal cortex and striatum and 2) motor cortex and striatum were identified and characterized based on automatically identified regions in the T2-weighted image and tractography results from DWI.

The MRI-based atlas of the mouse brain in (Ma et al., 2008) was considered for brain parcellation. To obtain more specific tracts, the sensory motor cortex defined in the original atlas was manually divided into sensory, frontal and retrosplenial cortex. The atlas template was elastically registered to each subject T2-weighted volume using ANTs (Avants et al., 2008), and the resulting transformation was applied to the label map to obtain the individual brain parcellation.

To estimate the fiber tracts, DWI was preprocessed, including eddy-current correction using FSL (Jenkinson et al., 2012), denoising (Coupe et al., 2008) and bias correction (Tustison et al., 2010). The five baseline images were averaged and registered to the T2-weighted image to correct for EPI distortions. Whole brain tractography was performed using a deterministic algorithm based on constrained spherical deconvolution model, considering as seed points the voxels where fractional anisotropy (FA)>0.1. The same threshold was defined as stop criterion for the algorithm.

The streamlines belonging to the two tracts of interest were selected based on the regions identified in the T2-w images. Frontal-striatum tract was defined as the streamlines with an extreme point in frontal cortex and passing through striatum. Motor-striatum tract was defined as the streamlines with a extreme point in motor cortex and passing through striatum. Once the fiber tracts were identified, its average FA and fiber density (FD) were computed. FD was defined as the number of streamlines normalized by its length and the regions’ volume as in (Batalle et al., 2014).

### Functional connectivity analysis

Two approaches were used to evaluate functional connectivity. On the one hand, whole-brain connectivity was evaluated using global network metrics (Rubinov and Sporns, 2010) and on the other hand seed-based analysis was performed to evaluate connectivity of frontal and motor cortex with the rest of the brain.

For both approaches, rs-fMRI was preprocessed, including slice timing, motion correction by spatial realignment using SPM8, correction of EPI distortion by elastic registration to the T2-weighted volume using ANTs (Avants et al., 2008), detrend, smoothing with a full-width half maximum (FWHM) of 0.6mm, frequency filtering of the time series between 0.01 and 0.1 Hz and regression by motion parameters. All these steps were performed using NiTime (http://nipy.org/nitime).

Region parcellation was registered from the T2-weighted volume to the preprocessed mean rs-fMRI. Whole-brain functional brain network was estimated considering the gray matter regions obtained by parcellation as the network nodes. Connectivity between each pair of regions was estimated as the Fisher-z transform of the correlation between average time series in each region. Network organization was quantified using graph metrics, namely, strength, global and local efficiency and average clustering coefficient (Rubinov and Sporns, 2010).

To perform the seed-based analysis, two regions -frontal cortex and motor cortex-were selected from the automatic parcellation. For each seed region, average time series in the seed was computed and correlated with each voxel time series, resulting in two correlation maps describing the connectivity of frontal or motor cortex with the rest of the brain. The following regions of interest were identified from brain parcellation to evaluate their connectivity with the seed region: striatum, globus pallidus, thalamus and cingulate, sensory motor cortices, frontal (if seed region was motor cortex) and motor (if seed region was frontal cortex). Connectivity was quantified as the mean value of the correlation map in each region, considering only positive correlations.

### Metabolite quantification

LC-Model (Provencher, 1993) was used to quantify the MRS-detectable metabolites, fitting a basis set of 19 small metabolites and 9 macromolecules. Quality criteria was defined for the whole spectrum as full width at half maximum (FWHM) lower than 0.06 ppm and signal-to-noise ratio higher than 5. In addition, metabolites quantified with relative Cramér-Rao lower bounds (CRLB) higher than 15% were discarded.

### GluCEST analysis

To assess glutamate from CEST acquisition, GluCEST contrast was measured as the asymmetry between the image obtained with saturation at 3ppm downfield from water (resonant frequency of exchangeable protons for glutamate) and the image with saturation at -3ppm as follows (Cai et al., 2013, 2012):

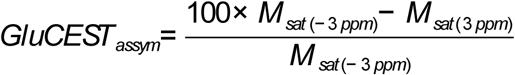

In order to quantify and compare GluCEST asymmetry, the ventricles were manually delineated in each GluCEST map and the minimum asymmetry value in this region was taken as a reference value to subtract from the whole-brain map. Average regional GluCEST asymmetry relative to ventricles was computed in manually delineated regions, namely, right and left striatum, and right and left cortex values were averaged.

### Stereotaxic surgery

Stereotaxic surgery was performed in 16-week old mice under isofluorane anesthesia (5 % induction, and 1.5 % maintenance). For optogenetic modulation, an adeno-associated virus (AAV) containing Channelrhodopsin (ChR2) under CamKII promotor was injected (AAV1-CamKIIa-hChR2(H134H)-eYFP-WPRE.hGH; AAV-ChR2) and a control YFP construct (AAV1-CamKIIa-eYFP-WPRE.hGH; AAV-YFP) was used as control. Virus production, amplification, and purification were performed by University of Pennsylvania-Penn Vector Core (titres: ∼1×10^12^ genomic particles/mL) and a volume of 0.5 μl of corresponding viral constructs were injected bilaterally in M2 cortex at the following coordinates: +2.46 anteroposterior (AP), ± 1 mediolateral (L) and -0.8 mm dorso-ventral (DV) from bregma and dura mater, using 5 μl Hamilton syringe with a 33 gauge needle at 0.1 μl/min. Injection needle was left for additional 5 min to allow diffusion of virus particles and avoid reflux.

For *in vivo* optogenetic stimulation, additional fiberoptic cannulas (MFC_200/240-0.22_3.5_ZF1.25_FLT; Doric Lenses) were placed bilaterally in the striatum at AP +0.14, L ± 2.2, DV -3 mm from bregma and duramatter and secured using dental cement. Optogenetic stimulation and behavioral testing was performed 4 weeks later.

For microdialysis experiments, AAV-YFP or AAV-CHR2 virus constructs were injected in the M2 as described above 4 weeks before microdialysis procedure. After 4 weeks, one concentric dialysis probe equipped with a Cuprophan membrane (1.5 mm long) was implanted in the dorsal striatum at coordinates (in mm, from bregma and skull): AP -1.1 (with a 20° angle); L 2.2; DV - 4.5 from dura matter, in anaesthetized mice (isofluorane, 5% induction and 1.5% maintenance). Additional bilateral fiberoptic cannulas were implanted as described above.

### Optogenetic stimulation

Blue light was delivered from 473 nm diode-pumped solid-state blue laser, (Laserglow) at 10 Hz, 5 ms pulse width and ∼5 mW (measured at the end of the patchcord) using a custom-made waveform generator (Arduino). For *in vivo* experiments, mice were habituated to be connected to the fiber optic patchcord (Doric Lenses) prior to the experiment. Zirconia sleeves (Doric Lenses) were used to connect the patchcord to the fiber-optic cannulas.

For electrophysiological recordings, fiber optic patchcord were placed above the slice surface.

### *In vivo* microdialysis

Extracellular glutamate (Glu) levels were measured by *in vivo* microdialysis as previously described (López-Gil et al., 2007). Microdialysis experiments were performed in freely moving mice 24 h (day 1) and 48 h (day 2) after surgery. The artificial cerebrospinal fluid (aCSF) was pumped at 1.65 µl/min and dialysate samples were collected every 6 min in microvials. Following an initial 3h stabilization period, six baseline samples were collected before 1 min optogenetic stimulation (Sample 7, day 1) and 5 min stimulation (Sample 7, day 2). Glutamate concentration was analyzed by HPLC consisting in a Waters 717plus autosampler and a Waters 600 quaternary gradient pump, and a Phenomenex Gemini 5μm 100×3mm column. Dialysate samples were precolumn derivatized with OPA reagent by adding 90 μl distilled water to the 10 μl dialysate sample and followed by the addition of 15 μl of the OPA reagent. After 2.5 min reaction, 80 μl of this mixture was injected into the column. Glutamate was detected with a Waters 2475 Fluorescence detector using excitation and emission wavelength of 360 and 450 nm respectively. Following sample collection, mice were sacrificed and brains were removed, sectioned and stained with neutral red to ensure proper probe placement.

### Multi-electrode arrays

Mouse brain sagittal sections were obtained on a vibratome (Microm HM 650 V, Thermo Scientific, Waltham, MA, USA) at 350 μm thickness in oxygenated (95% O_2_, 5% CO_2_) ice-cold artificial cerebrospinal fluid (aCSF) containing (in mM) 124 NaCl, 24 NaHCO_3_, 13 glucose, 5 HEPES, 2.5 KCl, 2.5 CaCl_2_, 1.2 NaH_2_PO_4_. 1.3 MgSO_4_, Slices were then transferred to oxygenated 32°C recovery solution of the following composition (in mM): 92 NMDG, 30 NaHCO_3_, 25 glucose, 20 HEPES, 10 MgSO_4_, 5 sodium ascorbate, 2.5 KCl, 1.2 NaH_2_PO_4_, 3 sodium pyruvate, 2 thiourea, and 0.5 CaCl_2_; for 15 min (Choi et al., 2019). Then, slices were transferred to oxygenated aCSF at room temperature and left for at least 1 h before recording.

Following recovery, slices were placed in a multi electrode array (MEA) recording dish and fully submerged in oxygenated aCSF at 37 °C. Electrophysiological data were recorded with a MEA set-up from Multi Channel Systems MCS GmbH (Reutlingen, Germany) composed of a 60 channels USB-MEA60-inv system with a blanking unit from Multi Channel Systems and a STG4004 current and voltage generator. Experiments were carried out with 60MEA200/30iR-ITO MEA dishes consisting of 60 planar electrodes arranged in an 8×8 array (200 μm distance between neighboring electrodes and an electrode diameter of 30 μm). Software for stimulation, recording and signal processing were MC Stimulus and MC Rack from Multi Channel Systems. Using a digital camera during recording assessed the position of the brain slices on the electrode field and the location of the laser for stimulation.

Striatal field post-synaptic currents (fPSC) were recorded in the dorsal striatum in response to stimulation of cortical afferences with short laser pulses of increasing intensity. Light stimulation was performed by placing the laser in the slice approximately 0.5 mm posterior to bregma, 2.5 mm ventral from surface, and 3 mm lateral from midline (Figure 3d-e, 6a). The evoked striatal fPSCs were analyzed after trains of 1 ms light stimuli of increasing intensity (0.00, 0.08, 1.5, 5.6, 13.7, 28.7, 47.7, and 60 mW/mm^2^).

For electrical stimulation one MEA electrode located approximately at 1 mm anterior to bregma, 2.5 mm depth from brain surface and 2,2 mm lateral from midline was set as stimulation one (Figure 6a, panel right), and fPSC were evoked by single monopolar biphasic pulses (negative/positive, 100 µs per phase). For input/output curves, cortico-striatal fibers were stimulated with trains of 3 positive-negative identical pulses at increasing currents (5 s inter-pulse time; 30 s train intervals; 250-3000 μV currents). Then, the pulse amplitude of subsequent stimuli was set to evoke 40% of the saturating fPSC response in the input/output curve. Long-term depression (LTD) was induced by theta-burst stimulation (TBS) consisting of 10 trains spaced 15 s apart. Each train consisted of 10 bursts at 10.5 Hz (theta), and each burst consisted of four stimuli at 50 Hz. Thus, the whole TBS stimulation period lasted 2.5 min (Hawes, et al. 2013). In all cases, the amplitude of evoked fPSC was quantified. Software for recording and signal processing was MC Rack from Multi Channel Systems. Using a digital camera during recording assessed the position of the brain slices on the electrode field.

### Behavioral assessment

All behavioral tests were performed during light phase and animals were habituated to the experimental room for at least 1 hour prior to testing. All apparatus were thoroughly cleaned between tests and animals. The order of the tests and time between tests is detailed in Figure 4d.

#### Open field

Spontaneous locomotor activity and exploratory behavior were assessed in an open field (OF) at week 20, 21 and 22 of age. The OF test was performed using a white square arena (40 × 40 x 30 cm) with dim light (∼20 lux). Mice were left in the center of the apparatus and were allowed to explore the arena during 11 min. Optogenetic stimulation was performed from minute 5 to 6 of the test. Animals were tracked and recorded with SMART 3.0 software (Panlab). Total number and time spent doing rearing and grooming events were measured to assess exploratory and stereotypic behavior.

#### Accelerating rotarod

Mice were placed on a motorized rod (30 mm diameter) with a rotation speed gradually increased from 4 to 40 rpm over the course of 5 min and the latency to fall was recorded to asses motor learning (Creus-Muncunill et al., 2019). The procedure was performed four times per day with 1h inter-trial interval for three consecutive days, 12 trials in total.

#### Balance beam

Motor coordination and balance was determined by measuring their ability to traverse a narrow beam according to (Creus-Muncunill et al., 2019). The beam consisted of a wooden square bar (50 cm long with 1.3 cm face), divided by 5 cm-frames and placed horizontally 50 cm above the bench surface, with each end mounted on a narrow support. Animals were allowed to walk for 2 min along the beam and number of frames crossed were measured.

### Golgi stain and dendritic spine quantification

Fresh brain hemispheres were processed using the Rapid GolgiStain Kit (FD Neurotechnologies), as previously described (Alvarez-Periel et al., 2017). Bright-field images Golgi-impregnated MSNs striatal neurons from 100 μm brain slices were captured using a Leica epifluorescence microscope (x63 oil objective, magnification 1,6). Image Z stacks were taken every 0.2 μm and analyzed with Fiji software. Dendritic segments were traced (>20-μm long; average, 47.35 μm; mean range, 20–95 μm) and the spines were counted. Spine density was calculated in three to thirteen dendrites per neuron, and the values were averaged to obtain neuronal averages (7-11 different neurons per animal, n=3 for WT-YFP, WT-ChR2 and HD-YFP and n=2 for HD-ChR2).

## Statistical analysis

All data are expressed as mean ± Standard Error of the Mean (SEM). Each point represents data from an individual mouse. Statistical analysis was performed using the two-tailed unpaired Student’s t-test or two-way ANOVA, and Bonferroni post hoc test as indicated in the figure legends. Values of p < 0.05 were considered as statistically significant.

## Data availability

All the data analyzed during this study is included in the manuscript and supporting files.

## Acknowledgments

We are very grateful to Ana López (María de Maeztu Unit of Excellence, Institute of Neurosciences, University of Barcelona, MDM-2017-0729, Ministry of Science, Innovation and Universities) and Maite Muñoz for their excellent technical support. This work has been funded by the Spanish Ministry of Sciences, Innovation and Universities through projects no. SAF2017-88076-R, RETICS de Terapia Celular, Instituto de Salud Carlos III (RD06/0010/0006) and la Marató de TV3. This research is part of NEUROPA. The NEUROPA Project has received funding from the European Union’s Horizon 2020 Research and Innovation Programme under Grant Agreement No. 863214

## Figure legends

**Supplementary Figure 1.**
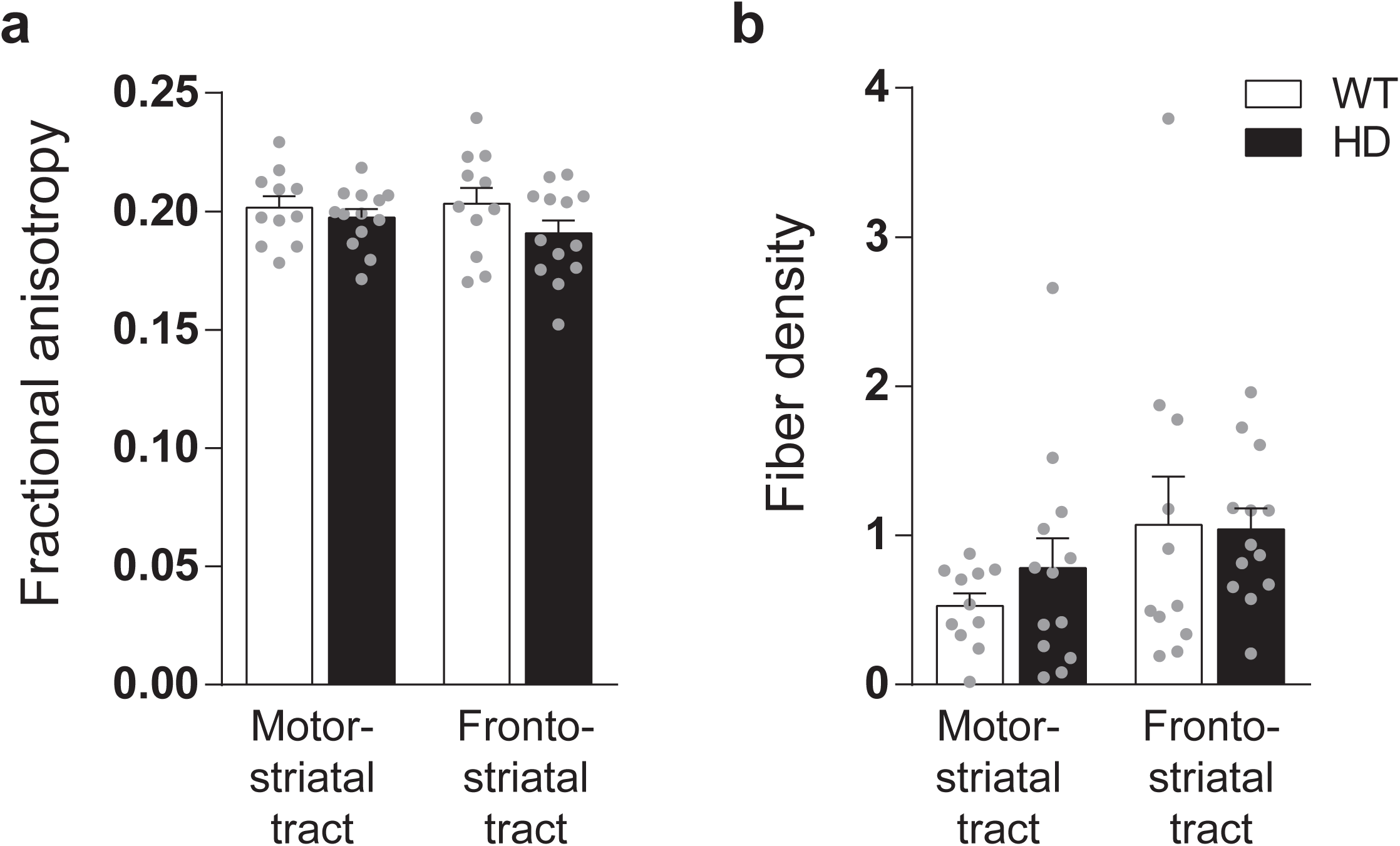
Fronto-striatal and motor-striatal diffusion and tract metrics are similar in WT and symptomatic HD mice. (**a**) Average fractional anisotropy and (**b**) fiber density of the fronto-striatal and motor-striatal fiber tracts estimated from DWI. Each grey point represents data from an individual mouse. Two-way ANOVA with Bonferroni’s post hoc comparisons test was performed. Data is represented as mean ± SEM (WT n=11 and HD n=13 mice). Each point represents data from an individual mouse.

**Supplementary Figure 2.**
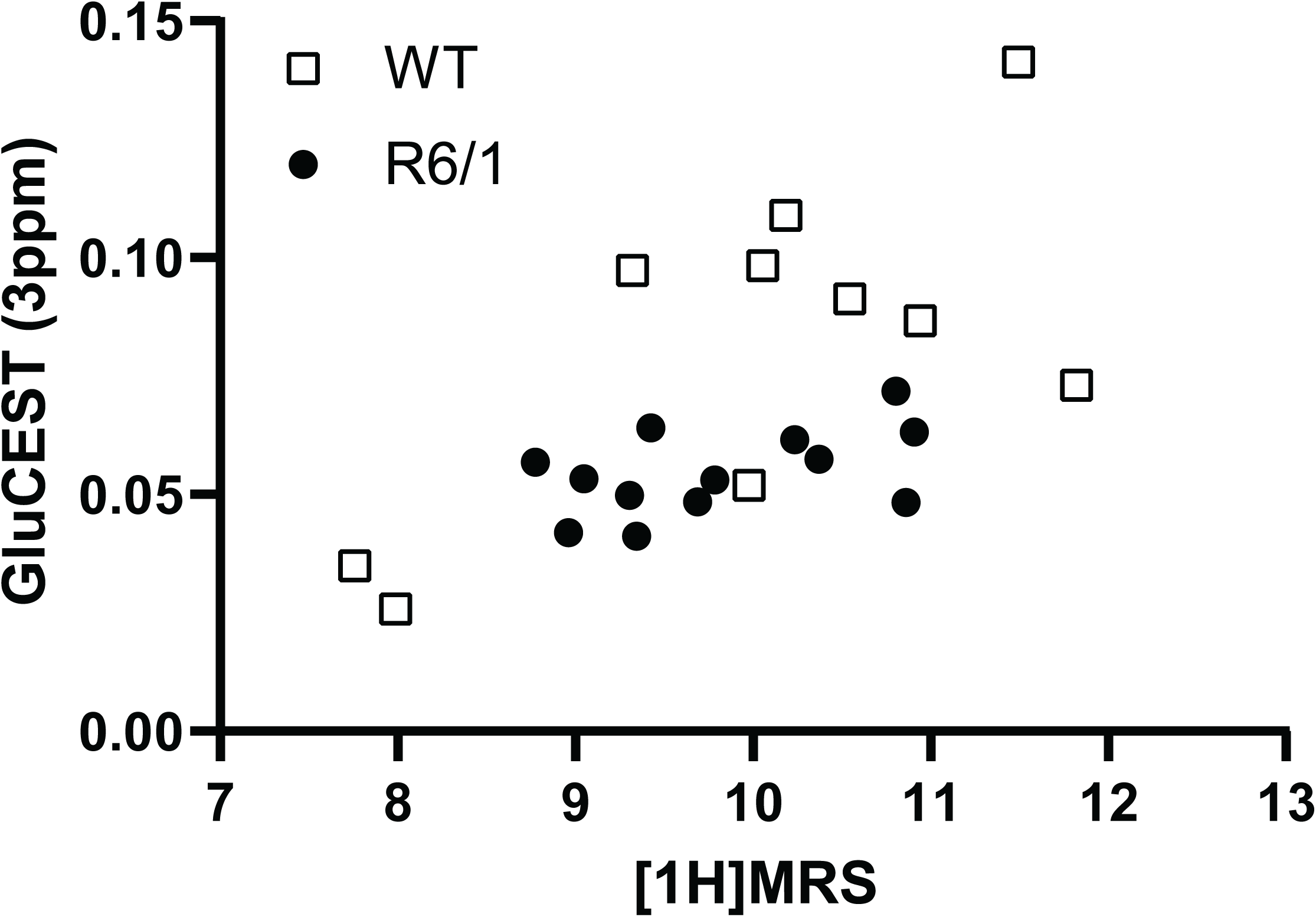
Correlation of glutamate levels obtained from GluCEST and 1H MRSin the striatum. Data is represented as mean ± SEM (WT n=10 and HD n=13 mice). Each point represents data from an individual mouse. Pearson correlation shows r=0.5979 with p=0.0026.

**Supplementary Figure 3.**
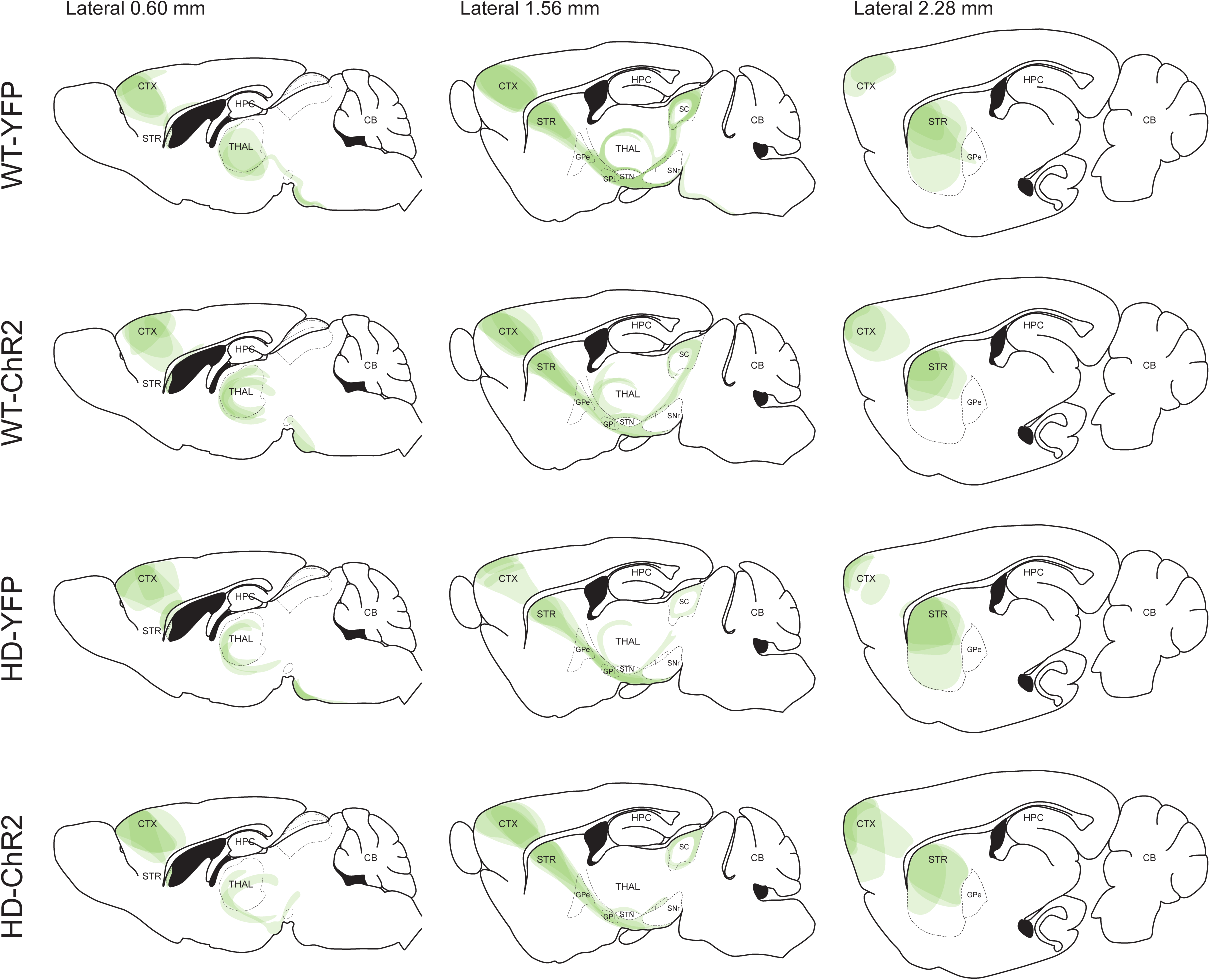
Validation of the AAV expression from M2 cortex and its projections by YFP fluorescence. Overlay of the YFP expression from different mice were drawn over three representative sagittal sections: medio-lateral coordinates 0.6, 1.56, 2.28 mmobtained from the brain atlas (Franklin and Paxinos, 1997). For each of the animals studied, brain slices were analyzed and the visualized virus-infected-pathways were plotted manually in the corresponding sagittal section. Each of the represented brain images contains different layers from four mice, each animal represented with a color opacity of 20%.

**Supplementary Figure 4.**
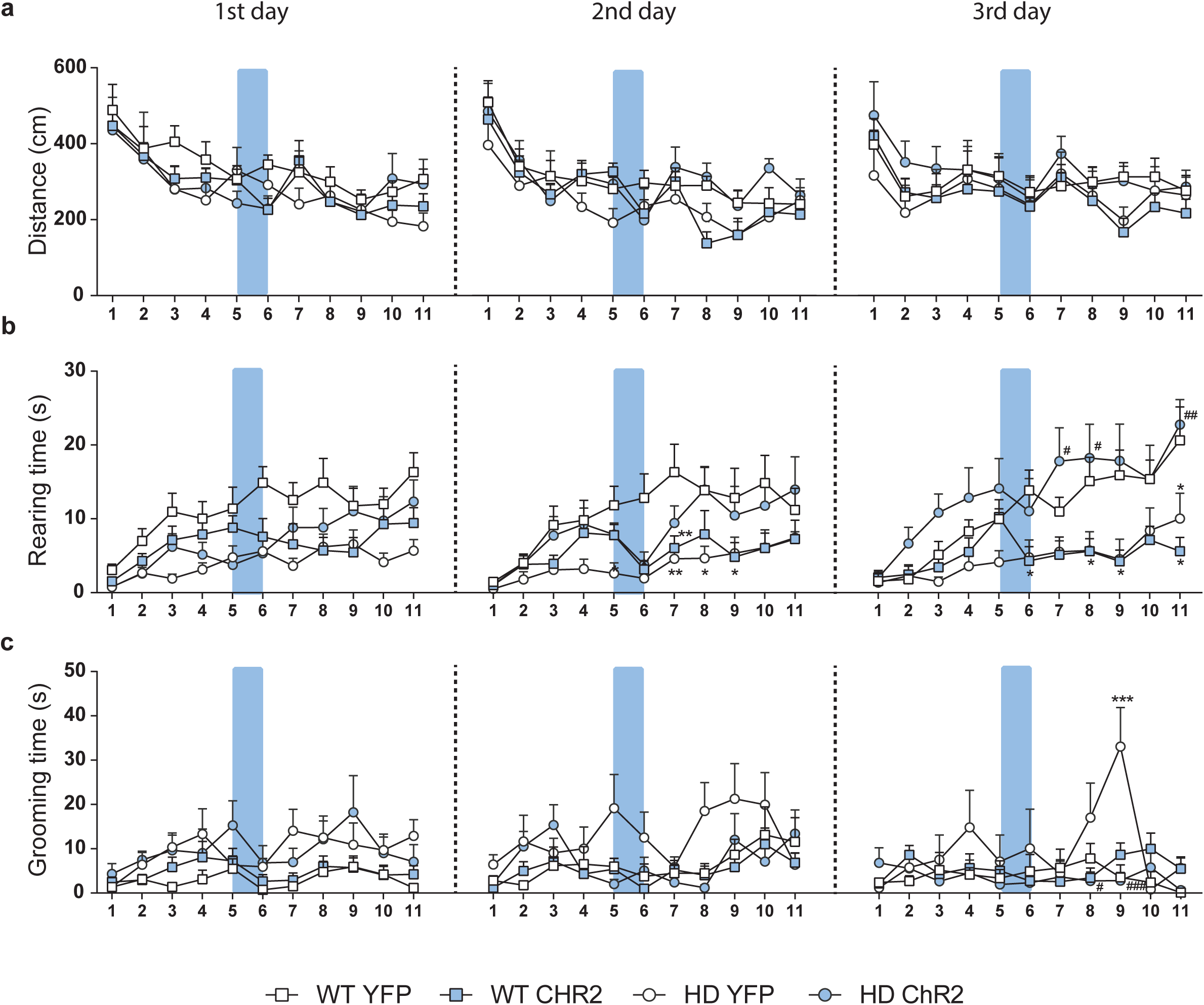
(**a**) Distance travelled (cm), (**b**) rearing time (s) and (**c**) grooming time (s) per minute in 20-week old WT-YFP, WT-ChR2, HD-YFP and HD-ChR2 mice during the first (left), second (middle) and third (right) open field sessions. Blue box represents the duration of blue light stimulation (1 min). Repeated measures ANOVA with group and time as factors followed by Bonferroni’s post hoc comparisons test was performed. Values are expressed as mean ± S.E.M (WT-YFP n=10, WT-ChR2 n=11, HD-YFP n=13 and HD-ChR2 n=11).

**Supplementary Figure 5.**
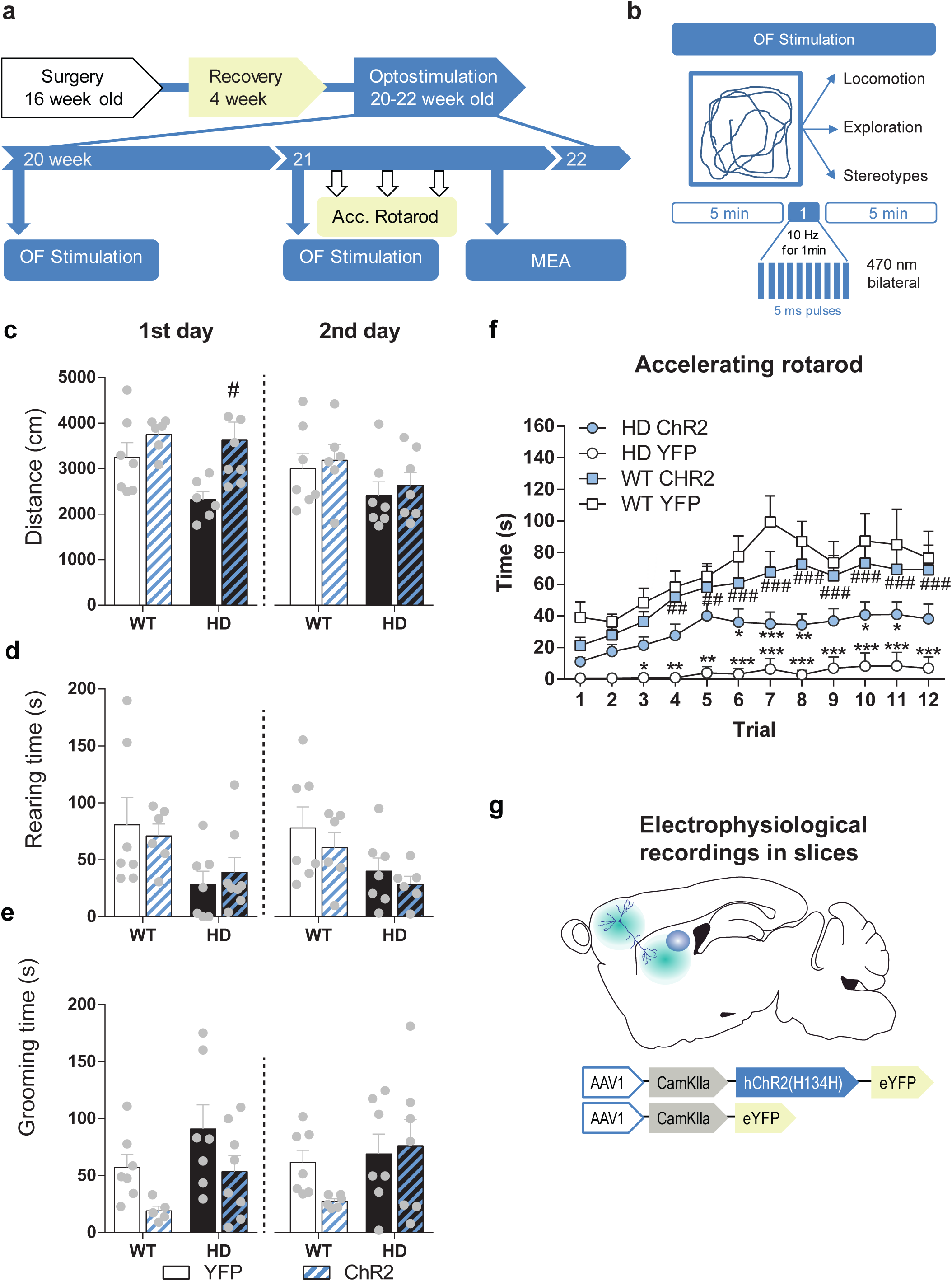
Behavioural data generated in mice used for figure 6a-d. (**a**) Surgery, behavior and optogenetic experimental timeline. The OF procedure was performed at 20 (1^st^ day OF) and 21 (2^nd^ day OF) week old mice. After 2^nd^ stimulation day, motor learning tests were performed and mice were sacrificed for electrophysiological analyses in brain slices. (**b**) Optogenetic stimulation consisted in 10 Hz light stimulation for 1 min during open field (OF) task and spontaneous activity was scored. (**c**) Locomotion (distance travelled), (**d**) exploration time (rearing time) and (**e**) stereotypes (grooming time) were evaluated during the first (left) and second (right) open field sessions. (**f**) Latency to fall in the accelerating rotarod task. (g) Schematic diagram showing AAV-ChR2 and control AAV-YFP constructs injections at M2 cortex and electrophysiological evaluation in sagittal cortico-striatal slices. Values are expressed as mean ± S.E.M (WT-YFP n=7, WT-ChR2 n=9, HD-YFP n=7 and HD-ChR2 n=8 mice). Each point represents data from an individual mouse. Data were analyzed by two-way ANOVA with genotype and light stimulation as factors, and Bonferroni test as a post hoc.

## Supplementary files

**Supplementary File 1**. Average seed-based BOLD correlation maps from WT mouse frontal cortex related to Figure 1a, visualized with ITK-SNAP software (Yushkevich et al., 2006).

**Supplementary File 2**. Average seed-based BOLD correlation maps from R6/1 mouse frontal cortex related to Figure 1a, visualized with ITK-SNAP software (Yushkevich et al., 2006).

**Supplementary File 3**. Average seed-based BOLD correlation maps from WT mouse motor cortex related to Figure 1a, visualized with ITK-SNAP software (Yushkevich et al., 2006).

**Supplementary File 4**. Average seed-based BOLD correlation maps from R6/1 mouse motor cortex related to Figure 1a, visualized with ITK-SNAP software (Yushkevich et al., 2006).

